# Hatchling fish disperse using an efficient multisensory strategy

**DOI:** 10.1101/2023.01.24.525385

**Authors:** Allia Lin, Efrén Álvarez-Salvado, Nikola Milicic, Nimish Pujara, David E. Ehrlich

## Abstract

Animals improve fitness by choosing when and where to disperse in the environment using sensory cues. In freshwater habitats subject to flood and drought, dispersal can urgently challenge newly hatched fish. Here we manipulated rearing environment and sensory systems to reveal an adaptive sensorimotor strategy for dispersal. If we constrained hatchlings or blocked feedback about motion by simultaneously impairing the lateral line and vision, they gulped air and elevated their buoyancy to passively sail on faster surface waters. In stagnant water, hatchlings then covered more ground with hyperstable swimming, tightly steering based on graviception. In hydrodynamic simulations, these adaptations nearly tripled diffusivity and made dispersal robust to local conditions. Through combined use of three senses, hatchlings adapt their behavior to flexibly and efficiently disperse.

## Introduction

Natal dispersal, movement from place of birth, allows organisms to track suitable environmental conditions, distribute genes, and collectively overcome local extinction events, ultimately impacting the robustness and evolution of populations [1]. Animals express their quality as dispersers and impact fitness by choosing when and where to move based on information about the environment [2, 3]. For some species, influencing dispersal presents acute challenges and opportunities during smaller, weaker juvenile stages, when external forces can dominate movement [4–8]. In these cases, animals that find ways to control dispersal can leverage external forces, mitigate risk while especially vulnerable, and dilute competition when kin can be most dense [9–11].

Precocious control over dispersal is particularly important for fish. Marine fish can use sensory cues to fight water currents and oppose dispersal, remaining in the safety of their home reef [12–14]. In contrast, freshwater fish can face urgency to disperse, as natal habitats quickly deteriorate, remodel, and fragment [15–17]. Capacity to sense the environment and control swimming in early development is best understood for zebrafish, native to the volatile waterways of the Indian subcontinent [18]. At the time of hatching, zebrafish are capable of evaluating the local environment through visual, lateral line (water flow), and vestibular (graviception and head acceleration) sensory modalities [19–23], but their small size and nascent nervous systems severely limit swimming ability [24, 25]. It remains unclear if hatchlings sense how well they disperse or use this information to adapt behavior.

We discovered hatchlings can detect conditions that impede dispersal and make adaptive responses to behavior and body. If we constrained hatchlings or impaired water flow and visual sensation, they elevated buoyancy by enlarging their swim bladders, a gas-filled organ they inflate by gulping air [26]. Consequently, they displaced to the surface where they flowed faster down a custom sluice mimicking natural waterways. Throughout one day of swimming under this multisensory impairment, hatchlings gradually swam more vigorously and engaged tighter feedback control to stabilize swimming horizontally, more efficiently covering ground. Labyrinthectomy revealed stabilizing feedback requires graviception, suggesting hatchlings regulate graviceptive control of swimming based on combined visual and flow sensation. We estimated the magnitude of effect of these mechanisms on dispersal using agent-based hydrodynamic simulations, finding distinct effects on key dispersal features: emigration and diffusion from siblings [27]. Hatchling adaptations nearly tripled diffusivity in stagnant conditions and primed their bodies for passive dispersal in flows. Together, these data show fish hatch with innate programs to sense environmental conditions and respond with adaptations that have an outsized impact on dispersal. We discuss underlying neural mechanisms and implications for population dynamics and fitness.

## Methods

### Animals

Four to eight days post-fertilization (d.p.f.), wild-type zebrafish of the AB strain were used for experiments unless otherwise indicated. Naturally spawned eggs were collected, cleaned, and maintained at 28°C in E3 solution (0.30 M NaCl, 0.01 M KCl, 0.03 M CaCl2, 0.02 M MgSO4, pH 7.1) on a 14/10 hour light/dark cycle at a density of 50 per 100-mm-diameter Petri dish. Larvae without inflated swim bladders at the time of manipulation or measurement were excluded from all experiments. Animal handling and experimental protocols were approved by the University of Wisconsin-Madison College of Letters and Sciences Institutional Animal Care and Use Committee.

### Behavior

Swimming kinematics were measured within the water column, adapted from a previous approach [28]. Briefly, six larvae from a single clutch were filmed in each glass tank (52×45×12 mm, G208,m Azzota Corp.) using a digital camera (GS3-U3-23S6M-C, FLIR) with a close-focusing, manual zoom lens (18-108 mm Macro Zoom 7000 Lens, Navitar). The field of view (approx. 2×2 cm) was aligned concentrically with the tank face. Transillumination was provided by a 5W, 940 nm infrared LED backlight (LED World) with aspheric condenser lens and diffuser (ACL5040-DG15-B, ThorLabs), and an infrared filter (43-953) removed reflections from an overhead, white LED strip (6500K, Adafruit). Occupancy as a function of depth quartile was measured in these tanks for 12 larvae, 10 minutes after acclimation, from lateral photographs with a 16 megapixel Sony camera (One-Plus). Four filming setups were mounted in parallel and enclosed within a light-tight box with an exhaust fan to cool to 28°C. Video was acquired at 40 Hz and the framewise position and orientation of larvae in the field-of-view was computed in real-time using the NI-IMAQ vision acquisition environment of Lab-VIEW (National Instruments). Swim bouts were detected with a 5 mm/sec speed threshold, bout duration was measured as the number of frames exceeding this threshold, and bout displacement was calculated as the Euclidian distance between positions at the start and end of this window using custom code in Matlab (Mathworks).

### Sluice

A custom fish sluice was prepared from a 27.7×1×1 inch (L×W×D) channel of 1/32 inch aluminum, friction fit with a trimmed, commercially available sluice mat designed for gold recovery, with alternating v-shape, square, and Hungarian riffles (ASR Outdoor). The sluice was installed at a 3° decline and room temperature aquarium water was flown down with a submersible pump (PE-1, Little Giant) connected to tubing with an open-jaw C-clamp (Humboldt) to constrain flow rate to 6.33-6.57 mL/sec. Anesthetized larvae were manually added at the top of the sluice with a transfer pipette and transit duration was timed with a stopwatch for 5 repetitions. If a larva took longer than 60 seconds to exit the sluice, which occurred on 8 of 117 trials, the trial was ended, the data point excluded, and the sluice flushed.

### Morphometry

Imaging and analysis for morphometry were performed by experimenters blinded to group. Bright-field photomicrographs of orthogonal dorsal and lateral perspectives of larvae were taken using a 12 megapixel iSight camera (Apple) through the ocular of a stereoscope (MZ12, Leica Microsystems). Larvae were immobilized dorsal up in 2% low-melting point agarose (Thermo Fisher Scientific 16520) and photographed before the agar was sliced vertically and rotated for a lateral-view micrograph. Profiles of the larvae and swim bladders were traced manually, and the dorsal- and lateral-view tracings were registered using custom MATLAB code, as previously [28]. We evaluated swim bladder morphology and fish density by fitting elliptic cylinders to morphometric measurements following a method adapted from Stewart and McHenry [29], assuming a swim bladder density of 1.31 mg/mL and tissue density of 1.04 g/mL, as previously [28].

### Confocal microscopy

Fluorescent hair cells in the *Tg(brn3c:mGFP)^s^*^356*t*^ line were imaged with a 60x water immersion objective (LUMPLFLN60XW, Olympus) on an inverted confocal microscope (FV1000, Olympus) [30]. Time-lapse images were acquired during ototoxin lesions by application of 50 *μ*M copper sulfate (Millipore Sigma) to the bath. Maximum intensity projections and depth-pseudocolored images were created in ImageJ [31].

### Lateral line lesion

Lesions were performed with 60 min soak of *Tg(brn3c:mGFP)* larvae at 5 d.p.f. in a 50 *μ*M solution of copper sulfate (MilliporeSigma) in E3, followed by 3 washes in E3. The lesion dose and timecourse was calibrated with live imaging using confocal microscopy. Swim bladder imaging began after 24 or 48 hour incubation in darkness following lesion. Lesions of animals used subsequently for behavior were confirmed by specific loss of neuromast fluorescence under a fluorescent stereoscope (SMZ-800N, Nikon). Sham-lesioned siblings underwent parallel soak in vehicle, wash steps, and fluorescence microscopy. Behavioral measurements were taken from 6 hatchlings of a single group per tank, starting 1 hour after termination of lesion and continued for 28 hours. Swimming kinematics were analyzed hourly and compared during blocks at the start to end of this period, comprising 9 hour windows before (start) and after (end) the dark phase of the entrained 14/10 hr light/dark cycle. Clutches were excluded from analysis if they contained insufficient data in either window (fewer than 50 bout distances or 3 tracked turn magnitudes for robust linear regression).

### Flow manipulation

Larvae were placed into 1 mL of E3 in a round-bottom, deep 96 well plate (V6771, Promega) in darkness in a 28°C incubator for 24 hours. Each well was 0.78 cm in diameter and 3.71 cm deep. A control group of 60 larvae per 100 mm Petri dish was incubated in darkness in parallel. Half of the well plates were incubated on an orbital shaker under constant, gentle rotation at 35 rpm (3520, Lab-Line).

### Bilateral labyrinthectomy

Larvae at 5 d.p.f. were anesthetized in 0.2% MS-222 (Syndel) in E3 for 5 minutes, then mounted on their side in 2% low-melting point agarose (Thermo Fisher Scientific 16520) and submerged in anesthetic solution. *Tg(brn3c:mGFP)* larvae were used to visualize vestibular hair cells and confirm lesion. The superficial otic vesicle was visualized on a widefield fluorescent microscope (BX51WI, Olympus) with 20x water immersion objective (XLUMPLFLN20XW, Olympus) under differential interference contrast. A microinjection needle (1.7 *μ*m diameter tip) was prepared on a micropipette puller (P-97, Sutter Instrument) from a borosilicate glass capillary (1B150F-4, World Precision Instruments) and backfilled with 1 mM copper sulfate (MilliporeSigma). For preliminary calibration, the injection solution was filtered (0.22 *μ*m filter, Millex) after incorporation of 128*μ*M sulforhodamine B (Sigma-Aldrich) to visualize the extent of diffusion under fluorescence. With visual guidance and control of a 3-axis micromanipulator (uMp, Sensapex), the pipette was used to puncture the otic vesicle en route to the proximity of the utricular otolith. Injection was produced with 1-5 pulses of 7 msec duration, 45 psi positive pressure to the micropipette, gated by an Openspritzer [32] triggered with custom code in LabVIEW (National Instruments). Injection was calibrated by sulforhodamine fluorescence filling the otic vesicle and later verified by comparable distension, and lesion quality and larval health were subsequently validated on an individual basis. After injection, the micropipette was removed, the larva freed from agarose and mounted on the contralateral side, and the protocol repeated for the second otic vesicle. Sham surgeries were performed by anesthetizing and mounting siblings identically, but piercing the otic vesicle with an empty and unpressurized microinjection needle. Larvae recovered overnight before visual verification of lesion by loss of green fluorescence in the otic vesicles.

Psychometric verification of labyrinthectomy was performed by mounting each larva in 2% low-melting point agarose (Thermo Fisher Scientific 16520) on the unmirrored side of a galvonometer (GVS011, Thor-Labs) and freeing its tail by removing agarose with a scalpel to film swimming. Swims were evoked with vibration of the galvonometer, comprising 5-cycle, 1 kHz sinusoidal motion of varying amplitude presented in descending order, with each amplitude delivered 5 times with a 5 sec inter-stimulus interval. Stimuli were composed in SutterPatch and commanded with D/A converter (dPatch, Sutter Instrument). The tail was illuminated with a 940 nm infrared, 48-LED array (Homyl) and was filmed with an infrared digital camera (BFS-U3-162SM, FLIR) with high-magnification lens (0.50x InfiniStix, Infinity Optical Company). Videos recorded at 400 fps were tracked with 5 equidistant points along the tail using a custom neural network for each larva, trained on 20 frames for 100,000 iterations using DeepLabCut [33, 34]. Following stimulus presentation, lesioned and sham larvae were unmounted and grouped into pairs for 48 hours of swim measurement based on similarity of stimulus sensitivity. Swimming was recorded as described above.

### Swimming simulation

To test how various manipulations affect hatchling movement in their habitat (primarily rivers, streams, ponds, and lakes), we consider an agent-based model in 2 canonical flow situations: (1) quiescent water mimicking a still lake and (2) open channel flow mimicking rivers and streams. Following standard models for microswimmer motion in flow [35–37], hatchling motion was modelled as a kinematic sum of the background fluid velocity (**u**), the hatchling swimming velocity (**v**_f_), and the hatchling settling velocity (**v**_s_).

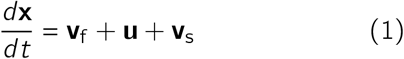

For quiescent fluid, the motion reduced to swimming and settling only. For open channel flow, we used the velocity profile calculated from direct numerical simulations of a zero-pressure-gradient flat plate boundary layer [38] that has been shown to match laboratory data of open channel flow [39]. Water depth was parameterized and constrained hatchling vertical position. The dimensionless velocity profile can be rescaled to various flow rates and water depths; here the ratio of friction velocity to the free stream flow velocity was chosen to be 0.05 based on experimental data [39]. This model captures the essence of the interactions between the hatchlings and ambient flow. We have neglected turbulent fluctuations and flow-induced reorientations based on observations that hatchlings actively maintain orientation (**Fig. 3**). Settling speed for larvae was calculated from empirical body densities (*ρ_p_*) following [40],

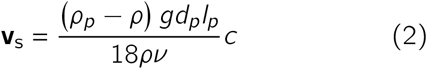

where *ρ* is the density of water at 28°C (996.3 kg*·*m^*−*3^), *ν* is its kinematic viscosity (1E-6 m^2^*·*sec^*−*1^), *g* is gravitational acceleration (9.81 m*·*sec^*−*2^), and *d_p_* and *l_p_* are the mean body diameter (3.17E-4 m) and length (3.90E-3 m), respectively, of the average hatchling. To calculate diameter, height and width were sampled along the length of each hatchling in our morphometric analysis and averaged. From [40], the shape factor was calculated to be *c* = 0.23 based on the measured aspect ratio (*a* = 12.31) and assuming, for simplicity, a horizontal orientation of the spheroid.

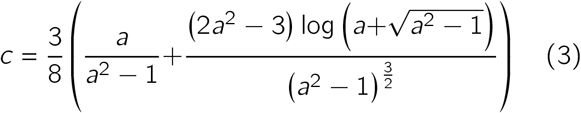

Each simulated hatchling had *ρ_p_* drawn randomly from empirical values. Eligible density values (in g/mL) for control hatchlings were [0.993, 0.983, 0.976, 0.975, 0.973, 0.973, 0.972, 0.964, 0.964, 0.961, 0.960, 0.957, 0.954, 0.952, 0.949, 0.938, 0.934, 0.926] and for flow-blind hatchlings were [1.011, 1.007, 1.006, 1.004, 1.001, 0.997, 0.994, 0.993, 0.992, 0.990, 0.989, 0.989, 0.988, 0.987, 0.987, 0.987, 0.986].

Larval swimming is modelled on empirical behavior, comprised of discrete bouts rather than continuous swimming. Between bouts, larval motion reduced to the background fluid velocity and settling velocity. The timing of swim bouts was governed by a Poisson process [28]. Larval position and orientation were calculated at 25 msec time intervals, the sampling frequency of empirical swimming data. Initiation of a bout occurred stochastically with a probability each time-step of 0.0435, the empirical probability of bout initiation per frame at 40 fps. Each swim bout comprised a fixed displacement applied evenly over 100 msec. Total bout displacement for larvae was typically 1.61 mm and for those adapted to low flow was 2.48 mm. To test the effect of swim bout displacement on sibling dispersal, we varied displacement and estimated the exponent (*ξ*) and coefficient (*k*) for the power law describing variance (*σ*^2^) as a function of time (*t*) across 5 simulations of 100 siblings each.

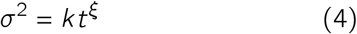

Where *ξ* = 1, this is a diffusive regime and we calculated the effective diffusivity (*D*) via *k* = 2*Dn*, where *n* is the number of dimensions in which the spreading is happening.

Larval orientation was defined in nautical convention as heading (angle projected on the horizontal plane) and elevation (angle relative to the horizontal plane) at each time-step. Larvae were initialized at random heading and with zero elevation. Simulated larvae never banked based on observations of a strong roll-righting reflex, keeping laterally level with respect to gravity [41]. To simplify implementation of constant bank angle, 3D turns while swimming were applied as extrinsic rotations to heading followed by intrinsic rotations in pitch (nose-up/down) that affected elevation but not heading or banking. This convention regularized simulations of swimming in 3D and in a vertical plane, with no impact on calculations of elevation when heading was held constant.

Turning behavior directed swim bouts. The direction of swimming velocity during a swim bout was defined by the heading and elevation of the larval body during that bout. Upon initiation of a bout, the larva generated rotations in heading and pitch. The rotation in heading was drawn from a Gaussian distribution (mean 0°, SD 25°) to approximate turns in a monolayer of water [42]. The rotation in pitch was repeated from the previous bout with 0.8 probability to simulate empirical hysteresis, and new rotations were randomly drawn from a Gaussian distribution matching empirical variance (mean 0°, standard deviation 3.80°). Swim bout heading and elevation were defined from larval orientation following these rotations and maintained throughout the duration of the bout. Alternatively, for simulations testing effects of density (Figure 1) but not larval control of orientation, the elevation of each swim bout’s velocity was instead randomly drawn from a Gaussian distribution with variance matching that of empirical swim bout elevation (mean 0°, SD 21.90°).

**Figure 1:**
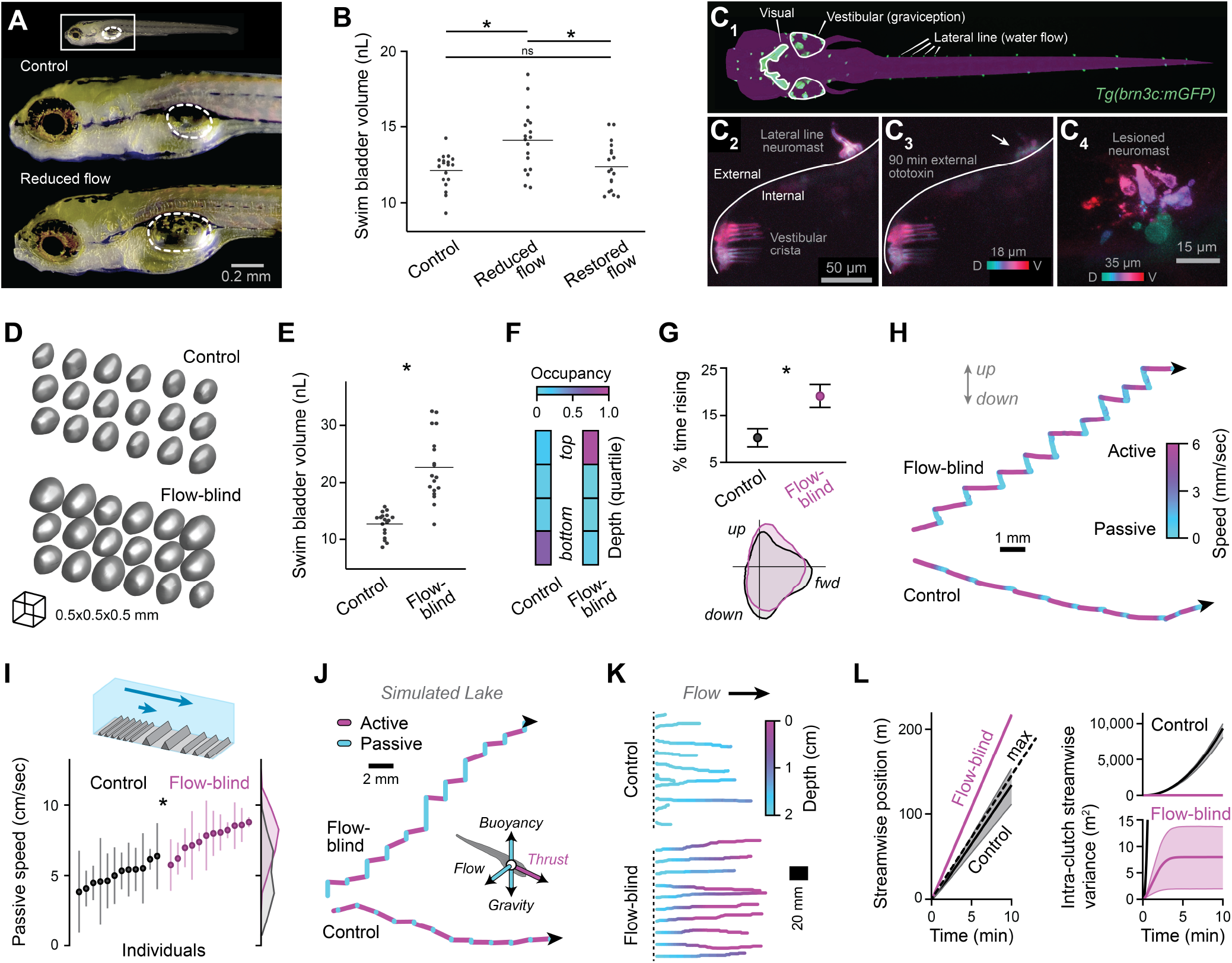
Hatchlings regulate dispersal via body density based on flow conditions. **(A)** Lateral perspective of a hatchling from low-flow environment and a control sibling, with the extent of the swim bladder outlined. **(B)** Swim bladder volume of hatchlings from low-flow environments, siblings from low-flow environments with flow artificially restored, and control siblings. One-way ANOVA with Bonferroni posttest (F_2,51_=7.85 *:p<0.05 and ns:not significant). **(C)** Hair cell responses to lateral line ablation. Dorsal-view, epifluorescent photomicrograph of hair cells of lateral line neuromasts and vestibular endorgans, as well as retinal ganglion cells (‘visual’) in a hatchling of the *brn3c:mGFP* transgenic line (C_1_). Confocal, depthcoded projection of a neuromast on the exterior of the otic vesicle and the posterior semicircular canal crista ampullaris interior to the otic vesicle, before (C_2_) and after (C_3_) 90 min. external copper treatment. Higher resolution, depth-coded photomicrograph of lesioned neuromast at 35 min. after copper exposure (C_4_). **(D,E)** 3D renderings of swim bladders (D) and measured volumes (E) from flow-blind hatchlings and control siblings 48 hr afer lesion (n=18). *p<0.05, two-way ANOVA (see Supp. Fig. S2), main effect of treatment: F_1,68_=80.8. **(F)** Occupancy of hatchlings in depths of tank by quartile (n=12). Z-test of occupancy at surface: z=3.69, p<0.05. **(G)** Proportion of time freely-swimming hatchlings moved upwards (*top*, within 45° of vertical), as mean and bootstrapped 95% CI. Polar probability distributions (*bottom*) of movement direction while passive (speed below 0.5 mm/sec). **(H)** Lateral views of swimming paths for one flow-blind hatchling (24 hours-post neuromast lesion) and one intact sibling, with instantaneous speed represented as color. **(I)** Mean flow speed in a custom sluice of anesthetized flow-blind and control hatchlings (n=12), as mean and standard deviation across 3-5 repetitions each. *p<0.05 by two-tailed t-test, t_22_=7.16. **(J)** Lateral views of simulated swimming paths with swim bladder volume of a flow-blind and control hatchling. Inset diagram shows forces acting on swimmers in simulation. **(K)** Representative paths (top-down) and depth (color-coded) for 10 simulated hatchlings with typical or flow-blind swim bladder volume, swimming for 5 sec in shallow, gently-flowing water (2 cm/sec). **(L)** Streamwise position and variance as a function of simulation time for clutches (100 hatchlings) with flow-blind or control swim bladder volumes, shown as mean and 95%CI. Dashed line indicates maximal passive transit at this flow speed.

Upon termination of a swim bout, after displacement was applied in the final time-step, the larva generated a pitch rotation towards horizontal. The magnitude of the rotation was drawn from a Gaussian distribution of mean *−wθ*, where *w* is the gain of the corrective rotation (0 for labyrinthectomized larvae, 0.126 for controls) and *θ* is the larva’s current elevation. The distribution had a standard deviation of 1.77°, calculated as the square root of the proportion of variance of empirical rotations while decelerating (+25 to +150 msec from bout peak speed) unexplained by pre-bout orientation [43].

### Data availability

All code, behavioral data, and morphometric data are available from the corresponding author upon request.

## Results

We first addressed whether hatchlings can respond to environmental features that impact dispersal. To test the hypothesis that hatchlings sense contexts that limit dispersal, we incubated individual, newly-hatched larvae in small (1mL) wells, as opposed to dishes full of siblings that agitate the water. After 24hrs, hatchlings in wells swam near the surface and had significantly enlarged swim bladders, organs they inflate by gulping air to elevate buoyancy (14.1 nL vs. 12.1 nL for controls, n=18; **Fig. 1A,B**) [26]. The specific feature of this context that stimulated inflation was lack of water flow, because hatchlings in wells exhibited no change in swim bladder volume if flow was maintained throughout incubation by gentle stirring (12.3 nL, n=18; **Fig. 1B**, ‘*restored flow*’). We conclude hatchlings can detect and respond to flow reduction by elevating their buoyancy.

To understand how hatchlings responded to chronic flow changes, we lesioned flow-sensitive neuromasts of the lateral line. These collections of superficial hair cells are deflected by flow and exposed to the water, permitting specific ablation by soaking in ototoxin [44–46]. In a transgenic line to visualize hair cells (*Tg:brn3c:mGFP*), we applied exterior copper sulfate to lesion the lateral line while sparing internal hair cells (**Fig. 1C, Supp. Fig. S1, Supp. Video 1**). These ‘flow-blind’ hatchlings nearly doubled the volume of their swim bladders at 24 hrs (23.2 vs. 13.8 nL for controls) and 48 hrs after lesion (22.7 vs. 12.7 nL), leading to a significant 2-4% reduction in estimated body density (**Fig. 1E, Supp. Fig. S2**; 24hrs: 0.964 vs. 0.992 g/mL; 48hrs: 0.961 vs. 0.995 g/mL). Consistent with elevated buoyancy, flow-blind hatchlings were most likely to be found near the surface (**Fig. 1F**; 92% vs. 17% for controls, n=12) and were more often rising when tracked swimming in depths of water (**Fig. 1G**, 7.56 concatenated hrs of tracked motion from 60 flow-blind hatchlings and 9.54 hrs from 66 controls). Flow-blind hatchlings moved along unique, staircase-like paths, thrusting forward with each bout of swimming, then rising while passive (**Fig. 1H**). When returned to normal conditions, flow-blind hatchlings survived well (89% vs. 93% for controls), recovered neuromast fluorescence consistent with lateral line regeneration [47], and exhibited typical swimming depth and swim bladder volume 6 days later (data not shown). Given that hatchlings responded similarly to low-flow conditions and ablation of the lateral line, we conclude they use the lateral line to detect flow reduction and respond with swim bladder inflation.

Hatchlings’ buoyancy response to low flow facilitated transport once flow was restored. We hypothesized the positive buoyancy of hatchlings that responded to low-flow environments would prime them to ‘sail’ on surface waters, where flow is fastest, if flow returned [48, 49]. To model natural transport in the laboratory, we flowed hatchlings down a gold sluice, a prospecting tool that mimics natural substrates where dense gold particles settle in rivers and streams. Flow-blind hatchlings flowed 50% faster down the sluice, even while anesthetized, confirming the hatchling response to low flow promoted passive transit once water flow was restored (**Fig. 1I**; 7.62 vs. 5.07 cm/sec, n=12). To test whether control of buoyancy affects dispersal on ecologically-relevant spatiotemporal scales, we made 3D hydrodynamic simulations of active and passive hatchling movements in natural flows. Hatchlings with larger swim bladders rose while passive (**Fig. 1J**), spent more time at the surface, and moved farther downstream (**Fig. 1K,1L, Supp. Fig. S3**). Typical hatchlings moved downstream slower than would a passive body at the surface, despite their active swimming, by taking refuge from flows at the substrate. In contrast, flow-blind hatchlings exceeded surface speeds by combining maximal passive movement with active swimming (**Fig. 1L**).

Hatchlings face a trade-off when deciding how to disperse, because buoyancy changes impede diffusion from siblings. We found that deviation from neutral buoyancy made transport more uniform across siblings (**Fig. 1K,1L**). Movement of typical siblings with approximately neutral buoyancy was superdiffusive, meaning they distributed faster than expected for a random walk (**Fig. 1L**). Distribution of siblings via swimming was compounded by differences in exposure to flow at different swimming depths, emphasizing the importance of natural variation of body density for dispersal (**Fig. 1B, Supp. Fig. S2B**). After hatchlings had elevated their buoyancy, however, their movement was subdiffusive, with uniform exposure to flow causing siblings to eventually move downstream in a cluster.

Given that hatchlings adapted their bodies to control dispersal, we next examined whether routine swimming reflected a propensity to disperse. In stagnant tanks, hatchlings swam strikingly horizontally (**Fig. 2A**). Horizontal swimming would promote active dispersal in natural habitats like lakes and streams with large horizontal span relative to depth. We previously showed hatchlings must actively swim to orient horizontally and counteract the destabilizing effects of flow and gravity [28]. By temporally aligning swim bouts, we found that hatchlings turned towards horizontal when they swam, specifically while they decelerated during each bout (**Fig. 2B**) [43]. Therefore, hatchlings actively work to stabilize their swimming horizontally.

**Figure 2:**
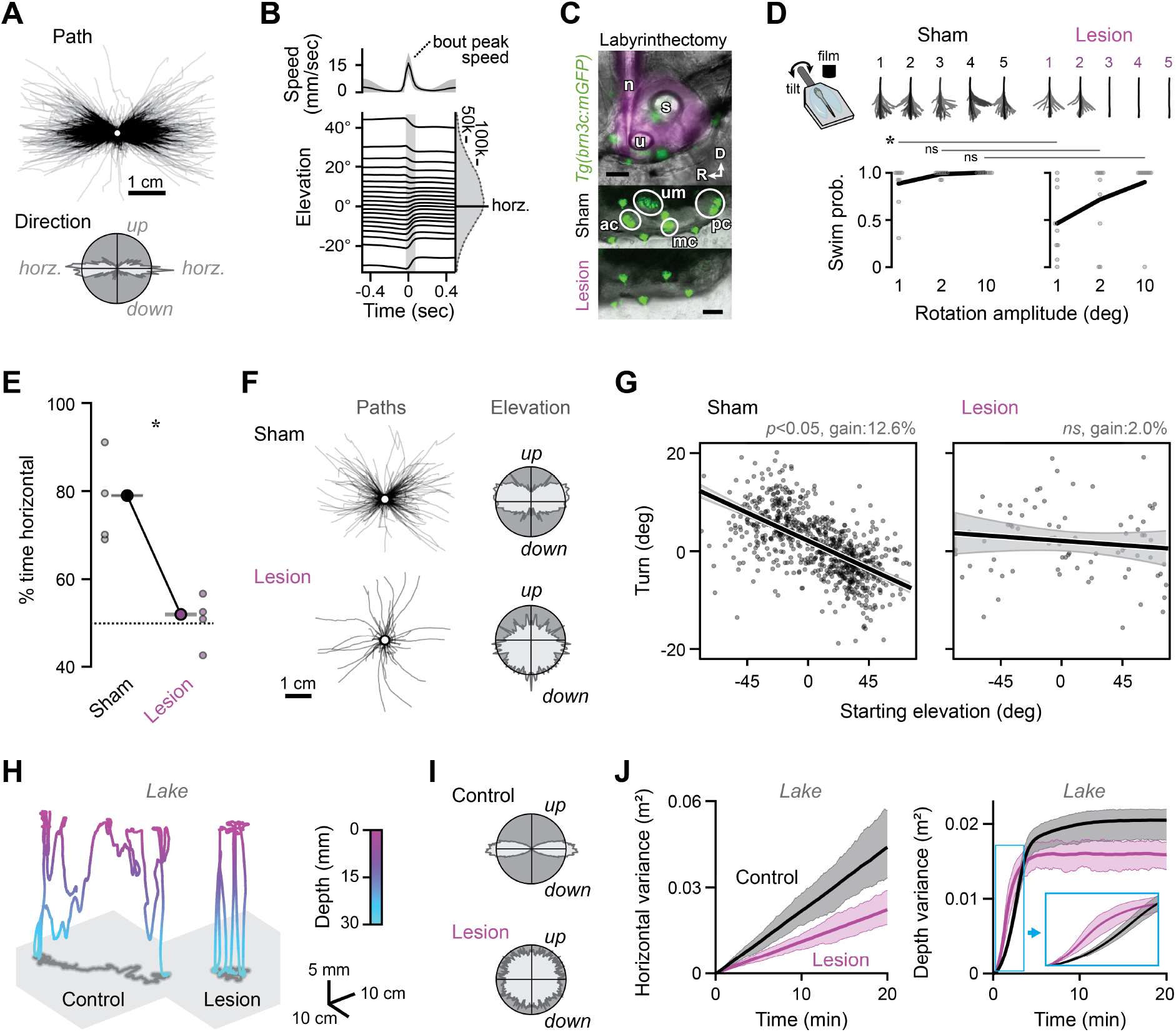
Hatchlings disperse by stabilizing swimming using graviception. **(A)** Swimming paths (*top*, n=1277) and trajectories (*bottom*, polar probability distribution) are biased horizontally. **(B)** Speed (*top*, mean *±* SD) and elevation (*bottom*, the body’s angle with the horizontal plane) during swim bouts aligned to peak speed. Bouts are sorted into 20 quantiles by pre-bout elevation, and the period of rotation towards horizontal is shaded. Marginal histogram counts frames at each orientation. **(C)** Labyrinthectomy via ototoxin pressure-injection through a micropipette into the otic vesicle, visualized with sulfrhodamine under epifluorescence (magenta, *top*). Fluorescent inner ear hair cells (*circles*, *Tg(brn3c:mGFP)*) are lost in lesioned but not sham-injected animals. Legend: ac-anterior crista, D-dorsal, mc-medial crista, n-needle, pc-posterior crista, R-rostral, s-saccule, u-utricle, um-utricular macula. Scale=50*μ*m. **(D)** Superimposed tracked tail segments in response to the first 5 rotational stimuli for representative sham and labyrinthectomized (lesion) hatchlings (*top*). Probability of swim response as a function of stimulus magnitude for individual hatchlings (points) and pooled means (lines). n=10, \**p*<0.05, *ns*: not significant by two-way ANOVA with Bonferroni post-test. **(E)** Proportion of time oriented horizontally (abs. elevation < 45°) during circadian day for groups of sham and labyrinthectomized hatchlings (small points) and pooled data with binomial SD (large points and bars). \**p*<0.05, z=117.5, for 50,898 pooled frames for lesioned hatchlings and 151,705 frames for controls. **(F)** Swimming paths and probability distributions of body elevation for sham and lesioned hatchlings. **(G)** For individual swim bouts, magnitude of the turn while decelerating (positive is upwards, negative downwards) is plotted as a function of elevation before the bout. Best-fit lines and 95% confidence intervals are plotted for bouts by sham (n=791) and labyrinthectomized hatchlings (n=81). **(H)** Simulated 3D swimming paths for one hatchling with an intact graviceptive system (12.6% turn towards horizontal each bout) and one with a labyrinthectomy (0% turn), color-coded by depth. **(I)** Polar probability distributions of body orientations of simulated hatchlings with 12.6% gain (”control”) vs. 0% gain (”lesion”). **(J)** Variance of horizontal (*left*) and vertical position (*right*) in time for 100 siblings in a 30 cm depth of lake as mean and 95% C.I.

We identified the sensorimotor mechanism by which hatchlings swim horizontally and promote dispersal. The orientation of flat bodies of water is reliably informed by the direction of gravity, perhaps more reliably than by light or water flow, and we hypothesized hatchlings steer towards horizontal using the ancient and rapidly-developing gravity sense [50–52]. To test this hypothesis, we developed a protocol for bilateral labyrinthectomy to acutely lesion the graviceptive vestibular system. Using this approach, we ablated vestibular hair cells while sparing superficial, flow-detecting hair cells of the lateral line, assessed visually in *Tg(brn3c:mGFP)* hatchlings (**Fig. 2C**). We validated each labyrinthectomy using psychometric testing based on swim response likelihood, finding impaired detection of weak vibratory stimuli (**Fig. 2D**) [53]. Afterwards, hatchlings swam normally in every respect but direction, moving and orienting isotropically (**Fig. 2E,2F**, **Supp. Fig. S4**). Lesioned hatchlings spent only 52.0% of their time oriented within 45°of horizontal, approximately at chance, compared to 89.0% for controls.

The graviceptive system stabilized horizontal swimming with high specificity. After lesion, turns during deceleration maintained a similar magnitude but were no longer conditioned on orientation relative to gravity (**Fig. 2G**). Whereas hatchlings typically turned 12.6% of the way back to horizontal with each swim bout (a stabilizing feedback gain of 0.126), turns were not correlated with orientation in labyrinthectomized hatchlings (gain of 0). The graviceptive system therefore provides a potent, intermittent feedback control mechanism to stabilize horizontal swimming. Control systems of this design elegantly accommodate steering maneuvers that change depth [43].

The graviceptive system proved critical for dispersal in low-flow conditions. When gravity-blind hatchlings were simulated in a lake, they failed to stabilize horizontally and covered less ground overall, despite swimming at the same intensity and rate (**Fig. 2H,2I**). Gravity-blind siblings exhibited half the effective diffusivity of intact siblings, distributing less horizontally (**Fig. 2J**). Additionally, intact siblings were slower to distribute throughout the water column but eventually adopted greater depth variance, as they tended to stratify and cover ground near the surface or substrate. Although hatchlings typically direct less of their swimming effort to changing depth than they did without this graviceptive response, they ultimately distributed in depth more effectively. Together, these data mechanistically link the ancient and rapidly-developing graviceptive system to dispersal at hatching.

Given the apparent role of swimming stability in dispersal, we tested whether hatchlings can modify stability based on sensory experience. By tracking behavior after loss of flow sensing, we found that hatchlings adjusted swim bout kinematics in parallel to their buoyancy. Specifically, hatchlings adapted by better stabilizing their swimming. By the end of incubation, flow-blind hatchlings in darkness had more than doubled the gain of their stabilizing turns (0.143 vs. 0.068, **Fig. 3A,3B**). With each swim bout, adapted hatchlings turned more than twice as far towards horizontal. Hatchlings also tended to increase gain following unimodal sensory loss, without either flow sensing (0.131 vs. 0.078) or vision (0.127 vs 0.100). In contrast, hatchlings failed to adjust their stabilizing gain if their senses were intact (0.084 vs. 0.071). Therefore, hatchlings adapt a stabilizing graviceptive behavior that promotes dispersal based on multimodal sensory feedback.

**Figure 3:**
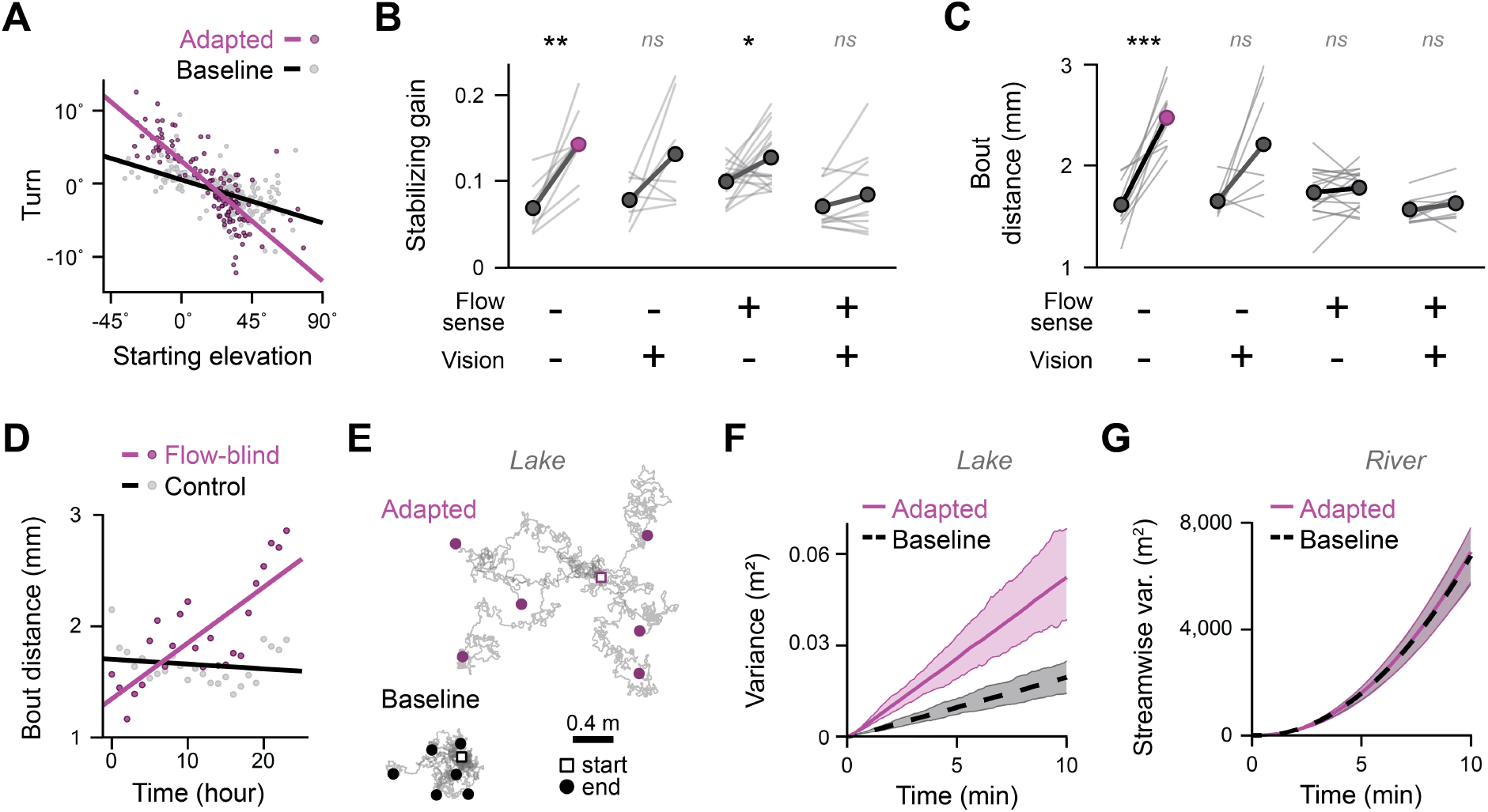
Hatchlings adapt swim intensity and stability to restore dispersal. **(A)** For discrete swim bouts from one representative clutch, magnitude of the turn while decelerating (positive is upwards, negative downwards) is plotted as a function of elevation before the bout. Bouts are compared at the start (*baseline*, n=189) and end (*adapted*, n=132) of swimming in darkness with lesioned lateral line. The slope of the best-fit lines reflect the stabilizing gain (baseline: 0.07, adapted: 0.18). **(B)** Stabilizing gain with and without flow and visual sensation (from lateral line lesion and darkness), plotted as paired values for each clutch at the start (left) and end (right) of 28 hrs of free swimming. Two-way ANOVA, main effect of flow sense: F_1,43_=9.9, *:*p*<0.05, **:*p*<0.005, *ns*:not significant by paired t-test with Bonferroni correction. Flow - vision -, n=10, t_9_=5.66; Flow – vision +, n=8, t_7_=2.12; Flow + vision -, n=18, t_1_7=2.85; Flow + vision +, n=11, t_10_=1.56. **(C)** Swim bout distance plotted as in B. ***:*p*<0.0005, *ns*:not significant by paired t-test with Bonferroni correction. Flow - vision -, t_9_=6.89; Flow - vision +, t_6_=2.56; Flow + vision -, t_17_=1.29; Flow + vision +, t_9_=2.09. **(D)** Hourly mean swim bout distance pooled for individuals in darkness in C, plotted with best-fit lines. R^2^ for flow-blind: 0.61, for control: 0.03. **(E)** Top-down projection of paths of 6 hatchlings with baseline swim bout distances (1.61 mm) and stabilizing gain (0.07) or those following adaptation (2.48 mm, 0.14 gain), simulated for 1 hour in a lake. **(F,G)** Variance of horizontal position in a lake (F) and streamwise position in a river (G) as a function of simulation time for clutches (100 hatchlings) with adapted or baseline swimming kinematics, shown as mean and 95%CI.

One possible explanation for swimming adaptation following loss of flow sensing and vision is that hatchlings combine sensory information from both modalities to evaluate motion levels. The lateral line conveys signals not only about external flow but also motion generated by swimming [54], providing a gross readout of transport. Given that zebrafish can intensify swimming under restraint in virtual reality [55], we next tested the hypothesis hatchlings swim more vigorously without lateral line or visual sensation. By the end of incubation under multisensory impairment, hatchlings generated bouts of swimming that covered a significantly greater distance (+54.3%, **Fig. 3C**) and lasted significantly longer (+24.2%, **Supp. Fig. S5A**). These effects accumulated gradually, with hatchlings swimming an extra 50 *μ*m per bout with each hour (**Fig. 3D**), and for an extra 1.7 msec per bout with each hour (**Fig. S5B**). In contrast, hatchlings with intact senses or unimodal sensory loss did not significantly adapt swimming (**Fig. 3C**), suggesting lateral line and visual inputs may provide a multisensory swim calibration signal.

To understand how these changes to swimming intensity and stability impact dispersal, we compared adapted and unadapted hatchlings in the simulated lake. Hatchlings with greater swimming intensity and stability covered more ground and distributed farther from siblings, nearly tripling their effective diffusivity (268% vs. unadapted swimmers, **Fig. 3E,3F**). These data confirm hatchlings can adapt swimming based on sensory feedback to dramatically improve dispersal, making more efficient use of each swim bout by better stabilizing horizontally.

If hatchlings are capable of swimming more intensely and stably, why would they engage this swimming mode conditionally? Simulations in the river provide an answer, as adapted swimming provided negligible benefit to movement downstream and dispersal from siblings (**Fig. 3G**). In fact, more vigorous swimming was counterproductive for distribution from siblings in flow, countermanding the advantages conferred by individual variation of density within a clutch (**Supp. Fig. S6**). Together, these data reveal hatchlings adapt swimming in the absence of sensory feedback confirming movement, reflective of low dispersal conditions. Therefore, hatchlings exhibit a tendency to make swimming more vigorous and stable in stagnant but not flowing water. Based on the observation that adapted swimming improves dispersal in stagnant conditions only, we conclude hatchlings modify their swimming to improve dispersal efficiently, not in flows where the energetic costs are unjustified.

## Discussion

Hatchling fish detected environmental conditions and adapted their bodies and behavior, revealing a surprising amount of control over dispersal that far exceeds the classical conception of hatchlings as plankton. Using a unique modeling approach incorporating detailed, empirically-derived kinematics and naturalistic flows, we evaluated how behavioral differences across individuals and states affected dispersal. Hatchlings made more efficient use of their limited swimming repertoire and nearly tripled their diffusivity by increasing swimming vigor and stability in stagnant conditions, revealing they can flexibly control dispersal through swimming. While the spatial scale of active contributions to swimming were relatively small, integration over hours and days would scale these distances to appropriate magnitudes to meaningfully impact ontogenetic habitat and therefore fitness [10, 56–58].

Consequences of active swimming adaptation were amplified through interactions with the background flow. Through small changes in buoyancy, hatchlings were able to maximize their passive transport and sail on flows, akin to how bubble-rafting snails passively cross subtropical oceans [59]. Variation in body density promoted diffusion *in silico*, revealing a potential downside to the sailing response: siblings efficiently emigrated but did so together. In ethological terms, the multisensory adaptations described would be particularly advantageous in the dynamic natural habitats, including ephemeral pools and oxbow lakes, of zebrafish and many freshwater species [18, 60]; if motion detection is diminished, for example in a constraining or evaporating pool that limits flow and swimming efficacy, the behavioral adaptations we observed may prepare hatchlings to maximize survival chances if new channels open or floodwaters arrive.

Sailing confers benefits for dispersal but may also have considerable costs. Buoyancy changes were only a temporary response to sensory changes, as hatchlings returned to normal density within days. It remains unclear if restoration of flow and regeneration of the lateral line was causative for recovery or if hatchlings would recover had sensation remained impaired. Consequences of positive buoyancy for predation risk or hunting success, both in terms of the kinematics of prey capture and the ability to track prey across depths, would be transient. Accordingly, hatchlings are not yet preoccupied with hunting owing to yolk stores, perhaps reflecting a critical period when dispersal is a higher priority than prey capture and, consequently, sailing is a viable strategy [11].

Zebrafish hatched with flexible behavioral programs and requisite neural pathways to control dispersal. Hatchlings efficiently promoted dispersal in stagnant conditions by stabilizing swimming using sensations of gravity for feedback control of orientation, specifically of elevation through rotations in pitch. They turned horizontally while they decelerated, potentially accommodating independent, serial generation of thrust [43]. Hatchlings set the gain of this feedback – how much each swim bout corrected deviations from horizontal – to hyperstabilize swimming based on multisensory visual and lateral line information, revealing an approach that not only improved dispersal in stagnant conditions but may generally afford flexible management of trade-offs between stability and maneuverability [61]. These data provide evidence for long-timescale modifications to swimming based on lateral line sensation that complement short-timescale responses during rheotaxis [62–64] and long-timescale hormonal changes [65]. Flow stimuli can elicit sustained activity changes in the serotonergic raphe [66], identifying a potential neural intermediate to transform prolonged flow changes into swimming adaptation and integrate flow and visual sensation. Consistently, serotonin can sensitize vestibular reflexes and serotonergic drugs can impact balance in humans [67, 68], and future tests will address if serotonin directly sensitizes graviceptive stabilization of swimming in hatchlings. Vestibular sensitization could in turn stimulate swim bladder inflation, which requires central vestibular activity, providing a common mechanism for adaptation [69].

The ability to control dispersal at hatching could have profound consequences for fitness and ecology of freshwater fish. Several examples of larval fish producing informed dispersal come from marine habitats, where older larvae than those studied here oppose dispersal using sensory cues [13, 14]. We found that larvae control dispersal earlier in development, within days of fertilization and hatching, and that freshwater fish actively promote dispersal instead of opposing it. Volatile habitats may select for behaviors that promote dispersal [70], and future studies will be required to evaluate how general these adaptations are across freshwater species or whether they correlate with habitat features. Precocious control over dispersal may confer resilience as habitat volatility increases and fish development accelerates with climate change [70–72]. The discovery that hatchling fish adapt to environmental conditions and elevate their chances of dispersal informs responses not only to natural habitat variation, but also anthropogenic disturbances to aquatic habitats [73, 74]. The dispersal strategy we describe for freshwater hatchlings comprises integration across sensory modalities to tailor behavior to local conditions and efficiently use a limited behavioral repertoire, revealing a mechanism for individuals to impact their own fitness and their species’ persistence.

## Acknowledgments

Research was supported by a gift from the Wisconsin Alumni Research Foundation and National Science Foundation grant CBET-2211704 to NP. We thank Han Wang for instrument access; Ashwin Bhandiwad, Jesse Weber, Kishore Kuchibhotla, and Tony Ives for helpful feedback; Joel Lord for fabrication; and Mary Halloran and David Schoppik for providing fish lines. This research was performed using the compute resources and assistance of the UW-Madison Center For High Throughput Computing (CHTC) in the Department of Computer Sciences. The CHTC is supported by UW-Madison, the Advanced Computing Initiative, the Wisconsin Alumni Research Foundation, the Wisconsin Institutes for Discovery, and the National Science Foundation, and is an active member of the OSG Consortium, which is supported by the National Science Foundation and the U.S. Department of Energy’s Office of Science.

## Author Contributions

Allia Lin: Conceptualization, Methodology, Investigation, Writing, Visualization. Efren Alvarez-Salvado: Conceptualization, Methodology, Investigation, Writing, Visualization. Nick Milicic: Investigation. Nimish Pujara: Conceptualization, Methodology, Software, Writing. David Ehrlich: Conceptualization, Methodology, Software, Investigation, Writing, Visualization, Supervision, Funding acquisition.

