## Supplemental Figures 1-6 for "Hatchling fish disperse using an efficient multisensory strategy"

### Supplementary Information

**Supplementary Figures**. Figures S1-S6 included below.

**Supplementary Movie 1**. Time-lapse movie of specific ablation of fluorescent lateral line hair cells (*Tg(brn3c:mGFP)*) in a neuromast on the exterior of the otic vesicle (top right) and in the posterior semi-circular canal crista ampullaris within the otic vesicle (bottom left). Time (in HH:MM) relative to external copper sulfate treatment. Voxels are color-coded according to depth. Gamma adjusted to 0.5 to visualize stereocilia.

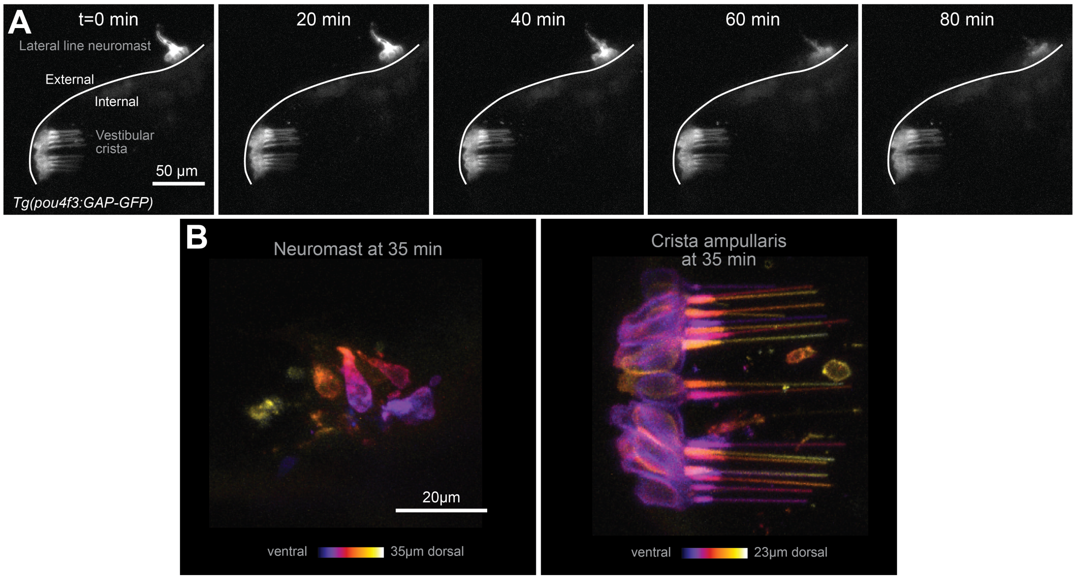


**Figure S1: Specific ototoxin ablation of lateral line hair cells. (A)** Time-lapse following external 50$\mu$M copper sulfate treatment of maximum projection photomicrographs of fluorescent hair cells (*Tg(brn3c:mGFP)*) in a neuromast on the exterior of the otic vesicle and in the posterior semicircular canal crista ampullaris within the otic vesicle. Gamma adjusted to 0.5 to visualize stereocilia. **(B)** Higher resolution photomicrographs of the neuromast (left) and crista (right) at 35 min post-treatment, color-coded for depth.

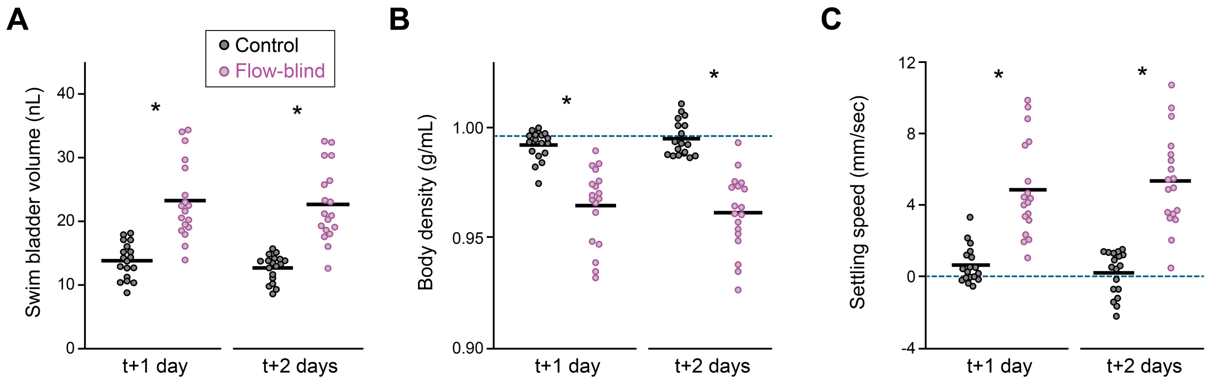


**Figure S2: Effects of lateral line lesion on body density and settling speed. (A)** Swim bladder volumes measured 1 and 2 days following lateral line lesion are shown for individuals (points, n=18) and group mean (bar), compared to unlesioned sibling controls. *$p<0.05$; Two-way ANOVA, main effect of treatment: $F_{1,68}=80.82$ ; not significant, main effect of day: $F_{1,68}=0.64$. **(B)** Body densities calculated from swim bladder volumes in A. Dashed line shows density of water at 28°C (0.996 g/mL). *$p<0.05$; Two-way ANOVA, main effect of treatment: $F_{1,68}=94.2$; not significant, main effect of day: $F_{1,68}=0$. **(C)** Settling speeds calculated from body densities in B. *$p<0.05$; Two-way ANOVA, main effect of treatment: $F_{1,68}=94.3$; not significant, main effect of day: $F_{1,68}=0$.

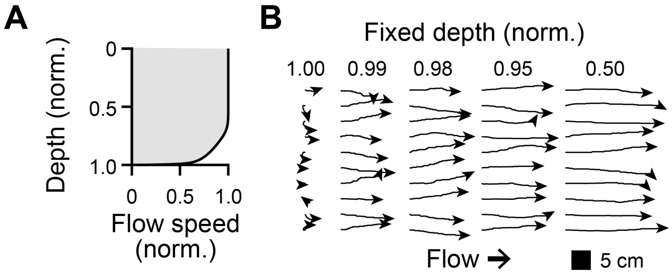


**Figure S3: Simulated flow and transport vary by depth. (A)** Flow speed as a function of depth used for simulations. **(B)** Example paths of 10 hatchlings simulated at fixed depth under 20 mm/sec flow.

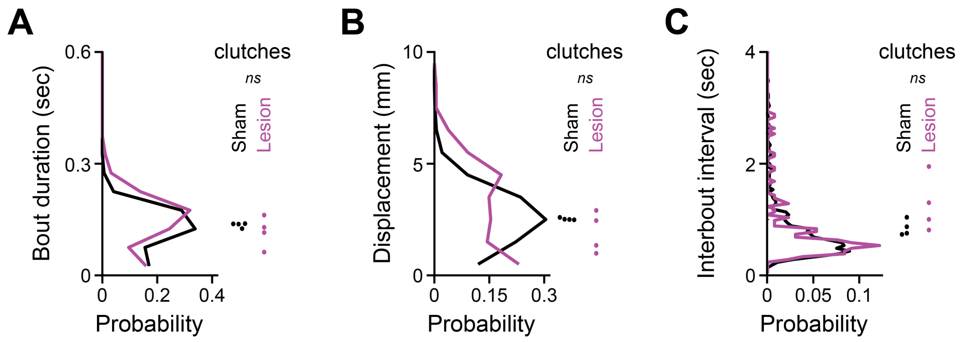


**Figure S4: Swim bouts are intact following labyrinthectomy. (A,B)** Probability distribution of swim bout duration (A) and displacement (B) for sham (1439 bouts) and lesioned hatchlings (209 bouts). Clutch-specific averages (n=4) are shown as dots. *ns*: not significant by two-tailed t-test. For bout duration, $t_6$=0.86; bout displacement: $t_6$=1.33. **(C)** Probability distribution of interbout intervals (start-to-start duration between successive bouts) for sham (995 interbouts) and lesioned hatchlings (131 interbouts). Clutch-specific averages (n=4) are shown as dots. Not significant, $t_6$=1.61.

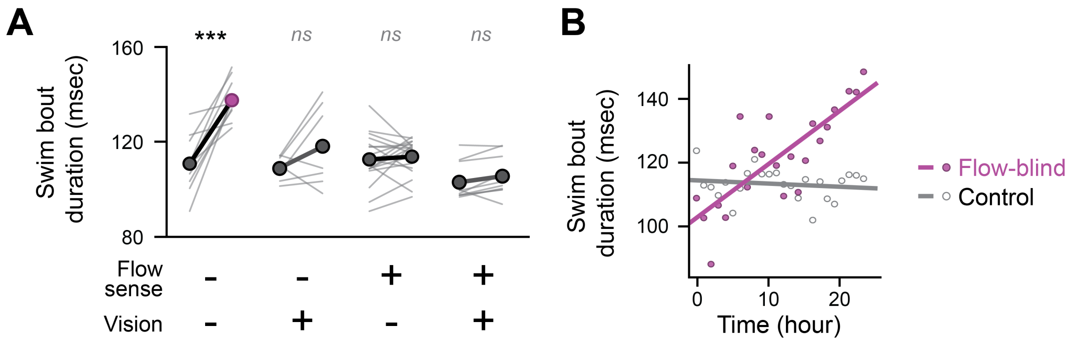


**Figure S5: Adaptation of swim bout duration under sensory impairment. (A)** Swim bout duration with and without flow and visual sensation (from lateral line lesion and darkness), plotted as paired values for each clutch at the start (left) and end (right) of 28 hrs of free swimming. ***:$p$<0.0005, *ns*:not significant by paired t-test with Bonferroni correction. Flow - vision -, n=10, $t_9$=6.08; Flow - vision +, n=7, $t_6$=1.86; Flow + vision -, n=18, $t_{17}$=0.46; Flow + vision +, n=10, $t_9$=2.71. **(B)** Hourly mean swim bout duration pooled for individuals in darkness in C, plotted with best-fit lines. $R^2$ for flow-blind: 0.64, for control: 0.02.

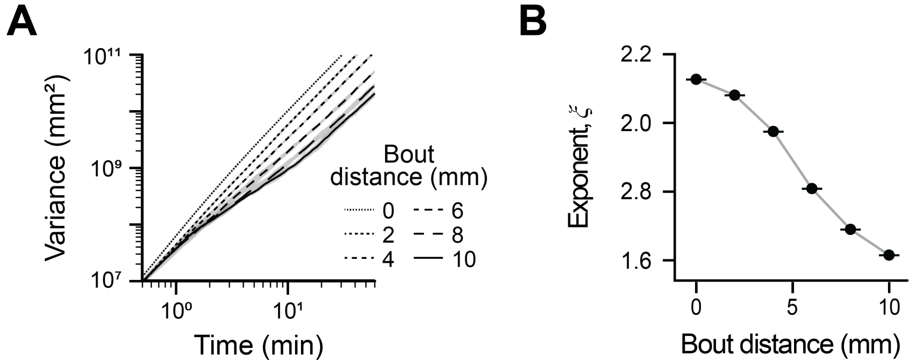


**Figure S6: Farther swim bouts reduce distribution from siblings in flow. (A)** Log-log plot of variance of siblings streamwise position vs. time, for 100 siblings across 5 simulations, plotted as mean $\pm$ standard deviation for 6 different bout distances. **(B)** Estimate of $\xi$ describing the superdiffusive relationships in A at each bout distance, from eq. 4, plotted as mean and standard deviation.
